# Cytosolic RNA binding of the mitochondrial TCA cycle enzyme malate dehydrogenase (MDH2)

**DOI:** 10.1101/2023.12.11.571136

**Authors:** Michelle Noble, Aindrila Chatterjee, Thileepan Sekaran, Thomas Schwarzl, Matthias W. Hentze

**Affiliations:** European Molecular Biology Laboratory (EMBL), Meyerhofstrasse 1, 69117, Heidelberg, Germany; Collaboration for joint PhD degree between EMBL and Heidelberg University, Faculty of Biosciences, Germany

**Keywords:** RNA-binding proteins, MDH2, metabolic enzymes

## Abstract

Several enzymes of intermediary metabolism have been identified to bind RNA in cells, with potential consequences for the bound RNAs and/or the enzyme. In this study, we investigate the RNA-binding activity of the mitochondrial enzyme malate dehydrogenase 2 (MDH2), which functions in the tricarboxylic acid (TCA) cycle and the malate-aspartate shuttle. We confirmed *in cellulo* RNA-binding of MDH2 using orthogonal biochemical assays and performed enhanced crosslinking and immunoprecipitation (eCLIP) to identify the cellular RNAs associated with endogenous MDH2. Surprisingly, MDH2 preferentially binds cytosolic over mitochondrial RNAs, although the latter are abundant in the milieu of the mature protein. Subcellular fractionation followed by RNA-binding assays revealed that MDH2-RNA interactions occur predominantly outside of mitochondria. We also found that a cytosolically-retained N-terminal deletion mutant of MDH2 is competent to bind RNA, indicating that mitochondrial targeting is dispensable for MDH2-RNA interactions. MDH2 RNA binding increased when cellular NAD^+^ levels (MDH2’s co-factor) was pharmacologically diminished, suggesting that the metabolic state of cells affects RNA binding. Taken together, our data implicate an as yet unidentified function of MDH2 binding RNA in the cytosol.

## Introduction

During the past decade, numerous metabolic enzymes at the heart of intermediary metabolism have emerged to bind RNA (Baltz et al., 2012; Castello et al., 2012; Ciesla, 2006; Hentze et al., 2018). Several metabolic enzymes (e.g., ACO1, GAPDH, TYMS, IMPDH, PKM2) have been found to moonlight in post-transcriptional regulatory functions (Casey et al., 1988; E. Chu et al., 1991; Constable et al., 1992; Dollenmaier & Weitz, 2003; Hentze & Argos, 1991; Hentze & Preiss, 2010; Kejiou et al., 2023; McGrew & Hedstrom, 2012; Nagy & Rigby, 1995; Singh & Green, 1993; Zhou et al., 2008). By contrast, the enzymatic activities of ENO1 and SHMT1 have been found to be riboregulated by cytosolic RNAs (Guiducci et al., 2019; Huppertz et al., 2022). While these examples underline physiologically significant roles of metabolic enzyme-RNA interactions, these cannot be assumed on the basis of binding data alone and the roles of most metabolic enzyme-RNA interactions await detailed exploration.

Malate Dehydrogenase has repeatedly been identified to interact with RNA across biological model systems ranging from *Escherichia coli* to human cells. Its RNA binding has been detected in multiple large-scale RNA interactome studies (Backlund et al., 2020; Baltz et al., 2012; Bao et al., 2018; Beckmann et al., 2015; Castello et al., 2012; Castello et al., 2016; Esmaillie et al., 2019; Garcia-Moreno et al., 2019; He et al., 2016; Huang et al., 2018; Kwon et al., 2013; Liao et al., 2016; Mallam et al., 2019; Marondedze et al., 2016; Matia-Gonzalez et al., 2015; Mullari et al., 2017; Panhale et al., 2019; Perez-Perri et al., 2023; Perez-Perri et al., 2018; Queiroz et al., 2019; Scherrer et al., 2010; Shchepachev et al., 2018; Trendel et al., 2019; Urdaneta et al., 2019; Wessels et al., 2016), purified together with polyA RNA as well as total RNA (Caudron-Herger et al., 2021; Gebauer et al., 2021; Schwarzl et al., 2022). Like most mitochondrial proteins, MDH2 is encoded by the nuclear genome, synthesised as a precursor protein in the cytosol, following which it is imported into the mitochondrial matrix (Chien et al., 1984; T. W. Chu et al., 1987; Grant et al., 1987; Grant et al., 1986; Sztul et al., 1988), where it assembles into functional homodimers. Together with its cytosolic isoform MDH1, MDH2 forms part of the malate aspartate shuttle, transporting reducing equivalents and metabolites across the mitochondrial membrane (Berg et al., 2002). It thus supports the preservation of the cellular redox state across compartments. Dysfunctions in MDH2 expression and activity are involved in several pathologies, including different forms of cancer and neurodevelopmental disorders (Ait-El-Mkadem et al., 2017; Cascon et al., 2015; Liu et al., 2013; Lo et al., 2015).

MDH2 belongs to the 2-hydroxy acid dehydrogenase superfamily and catalyses the reversible conversion of malate to oxaloacetate with the simultaneous reduction of NAD^+^ to NADH. Like other dehydrogenases, MDH2 has an NAD^+^-binding Rossman fold, which has been proposed as an RNA-binding domain (Castello et al., 2015; Hentze, 1994; Liao et al., 2016; Nagy et al., 2000). MDH2’s RNA-binding function has been implicated in the post-transcriptional regulation of SCN1A mRNA by binding to its 3’UTR (Chen et al., 2017). It has also been reported to promote tumorigenesis via lncRNA AC020978 (Xu et al., 2021). Additionally, MDH2 binding to lncRNA GAS5 was linked to TCA cycle regulation by disrupting the formation of a MDH2-FH-CS metabolon complex (fumarate hydratase: FH, citrate synthase: CS) (Sang et al., 2021). Although these studies uncover possible functions of the RNA-binding activity of MDH2, they have focused on individual MDH2-RNA interactions and an unbiased, comprehensive analysis of MDH2’s RNA-binding properties is still lacking.

We therefore employed eCLIP (Van Nostrand et al., 2016) to determine the scope of RNA binding by MDH2. In addition, we provide insights into the cellular locale of MDH2-RNA interactions as well potential effects of MDH2’s cofactor NAD^+^ on its apparent RNA-binding activity.

## Results

### Human mitochondrial malate dehydrogenase (MDH2) binds RNA in Huh7 cells

We probed the RNA binding of MDH2 in the human hepatocarcinoma cell line Huh7 using two orthogonal biochemical approaches. For the polynucleotide kinase (PNK) assay (Richardson, 1965), cells were exposed to ultraviolet light (UV 254 nm, 150 mJ/cm^2^) to induce covalent bonds between RNA-binding proteins and their bound RNAs, followed by MDH2 immunoprecipitation (IP) from cellular lysates. The 5’ ends of MDH2-associated RNAs were subsequently labelled *in vitro* with γ-^32^P ATP, and assessed by denaturing gel electrophoresis. In the PNK assay for MDH2, RNase-sensitive autoradiography signals were detected after MDH2 immunoprecipitation. When the RNase digestion is limited, large MDH2-RNA complexes migrate as a smear on a gel, indicating diverse lengths of the RNAs bound to MDH2. Following more extensive RNase digestion, the signals condense to a band slightly above the expected molecular mass of native MDH2. An identical assay with isotype matched IgG as a specificity control showed only minor background signals. (Fig. 1A). Moreover, we also find that the RNA signal is dependent on UV-crosslinking, further confirming its specificity (Supplemental Fig. 1).

**Figure 1.**
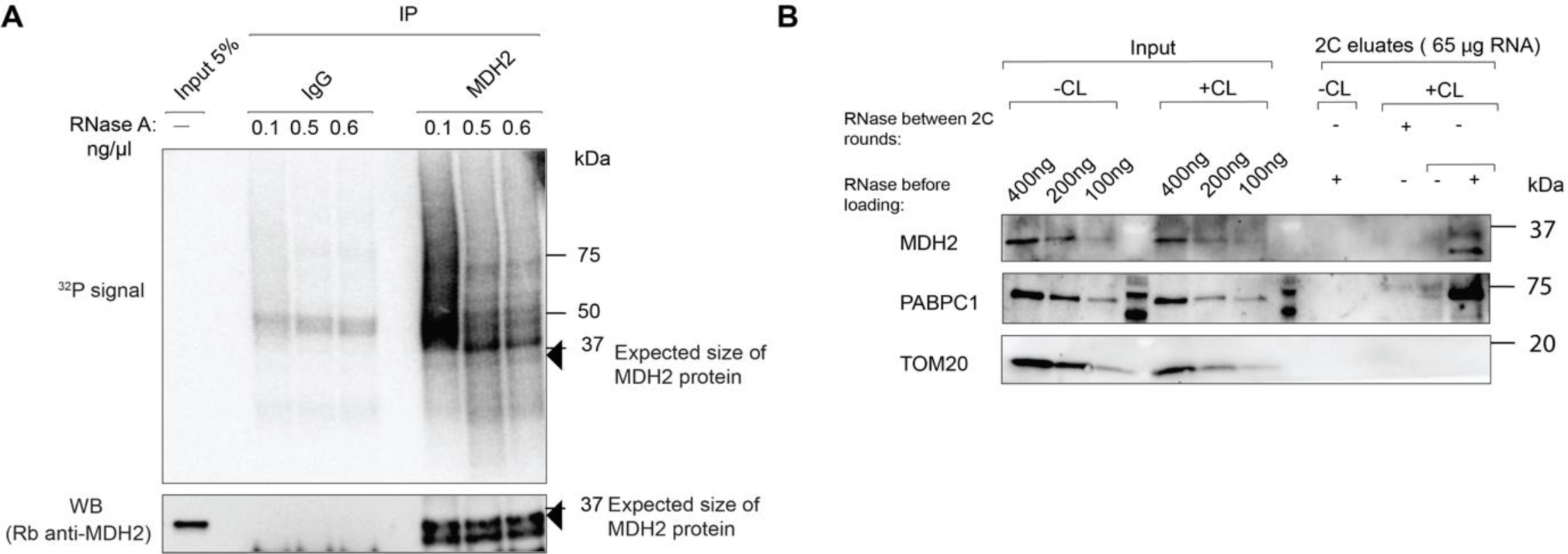
MDH2 binds RNA in cells. (A) RNA-binding activity of MDH2 in PNK assay. Top panel shows the phosphor image to visualise RNA co-purified with MDH2 following γ-32P ATP labelling by polynucleotide kinase (PNK). Bottom panel shows a representative western blot to confirm equal IP efficiencies. Black arrows indicate the expected molecular mass of MDH2. (B) RNA-binding activity of MDH2 assessed by 2C assay. Specific MDH2 western blot signals are detected when the sample has been treated with RNase before loading to visualise it around the expected molecular mass. RNase treatment before a second round of 2C abolishes the western blot signal. PABPC1 was used as a positive control and TOM20 as a negative control.

The complex capture (2C) assay identifies RNA binding based on RNA-mediated protein retention on silica columns that are routinely used for RNA purification (Asencio et al., 2018). Following UV-crosslinking, samples eluted from the silica column are assessed for the presence of proteins of interest by western blotting. Before gel analysis, the eluates are digested with RNase to minimise the contribution of RNA to the molecular mass of the RNA-protein complex and to visualise the protein near its expected molecular mass. 2C reveals that MDH2 is retained on the silica column in a UV-crosslinking dependent way and requires RNA, because RNase treatment prior to a second round of 2C abolishes MDH2 retention. Canonical RNA-binding proteins like PABPC1 (polyA binding protein) show a similar behaviour, while TOM20 (without identified RNA-binding activity) used as a specificity control is not co-purified (Fig. 1B).

### MDH2 binds predominantly cytosolic RNAs

Having confirmed cellular RNA binding of MDH2, we used enhanced crosslinking and immunoprecipitation (eCLIP) (Van Nostrand et al., 2016) to determine which RNAs are bound by MDH2. RNAs crosslinked to MDH2 were identified by sequencing after MDH2 immunoprecipitation (IP) from total cell lysates rather than lysates from purified mitochondria to avoid a data bias based on a-priori assumptions about where MDH2 may bind RNA. A control immunoprecipitation experiment performed with isotype matched IgG was also used to determine binding specificity.

After cDNA preparation, we ensured the quality of the sequencing libraries by optimising the number of PCR cycles required for library preparation in test PCRs (with 10% of the cDNA samples) followed by PAGE analysis. The final replicate libraries were pooled and assessed using their bioanalyzer profiles (Supplemental Fig. 2A). We generated three independent replicate sequencing libraries for MDH2 IP (∼9.1 million uniquely mapped reads/sample), SMI (∼4.3 million uniquely mapped reads/sample) and isotype-matched IgG IP (∼4.3 million uniquely mapped reads/sample). Principal component analysis confirmed close clustering of respective replicates distinct from the clusters of other samples (SMI/MDH2 IP/IgG IP) (Supplemental Fig. 2B)

In total, we identified 524 distinct, specifically enriched binding regions for MDH2 (Supplemental Table 1) across 361 unique RNAs, a major proportion of them on RNAs expressed outside of mitochondria (Fig. 2A shows the distribution of crosslinking counts across MDH2 targets). This observation was unexpected, considering MDH2’s primary enzymatic function and its enrichment in the mitochondrial matrix (Aziz et al., 1981; Musrati et al., 1998; Sztul et al., 1988). MDH2’s unexpected preference for cytosolic RNAs (transcribed from nuclear genomic DNA) over the abundant MT-RNAs (transcribed from mitochondrial DNA) concentrated in the locale of the mature protein suggests that its RNA association may primarily be relevant in the cytosol.

**Figure 2.**
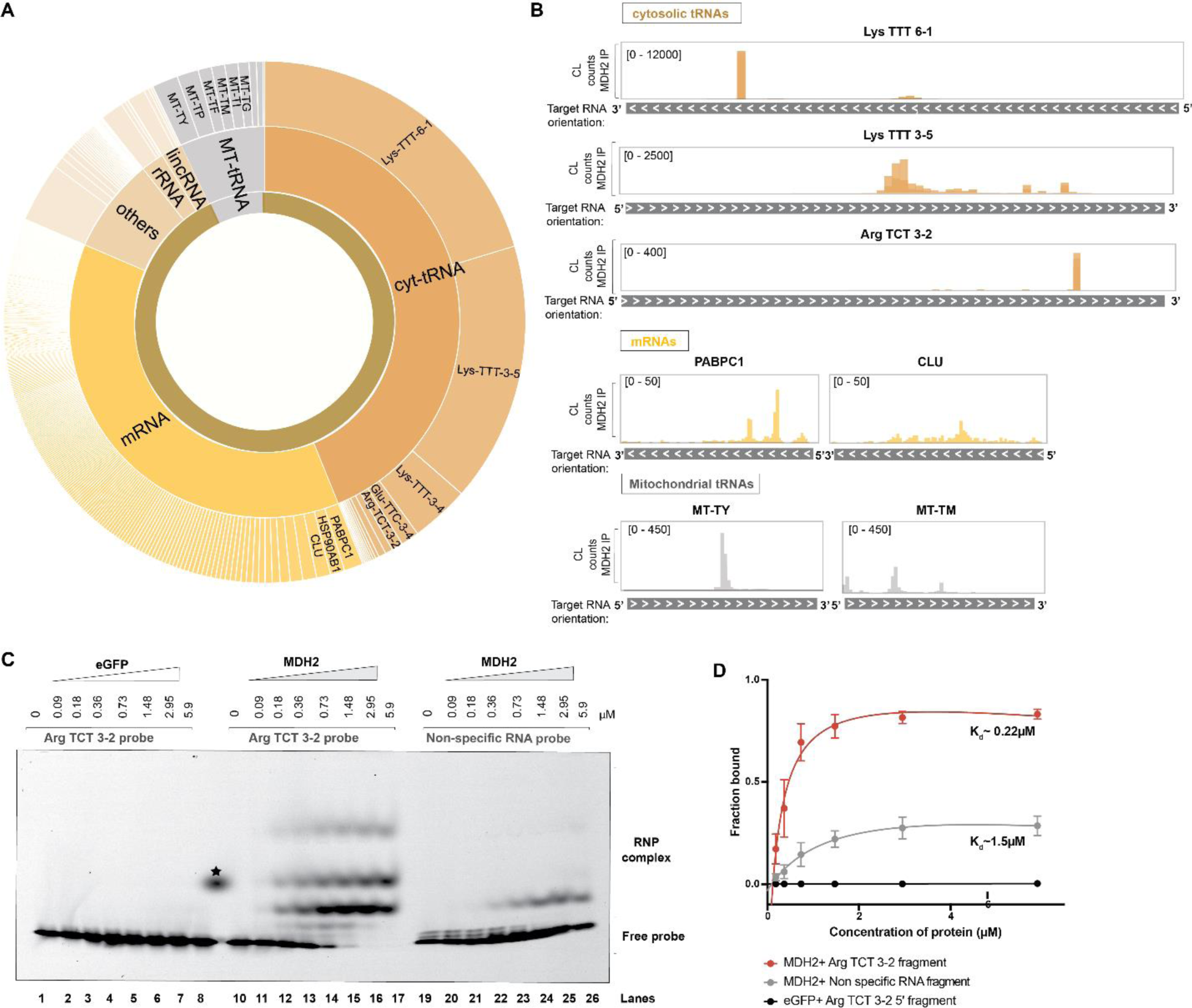
MDH2 predominantly binds cytosolic RNAs. (A) Sunburst chart showing the distribution of crosslinking counts in binding sites identified from MDH2 eCLIP data. “Cytosolic” indicates nuclear-encoded RNAs and “Mito” indicates mitochondrially-transcribed RNAs. (B) Crosslinking (CL) count distribution of selected target RNAs bound by MDH2 visualized using the IGV interactive tool (Robinson et al., 2011). (C) Representative EMSA assays performed to assess the affinity of MDH2 to a tRNA Arg TCT 3-2 probe or a control RNA probe (RPS23 5’ UTR sequence). Binding of eGFP to the target RNA probe is included as a negative control. (D) Binding affinities from EMSA experiments were estimated using a non-linear regression fit (solid line). Data reported as mean ± S.D (dots with error bars). Number of experiments, n=3-4 from two independent protein purifications.

Among the mitochondrial RNAs associated with MDH2, a preference of MDH2 for tRNAs emerged. MDH2 binds 9 of the 22 MT-tRNAs (Suzuki et al., 2011) expressed in human mitochondria (Supplemental table 1). MDH2’s propensity to bind tRNAs is also reflected amongst the cytosolic RNA interactors, with cytosolic tRNAs (specifically Lys TTT isoacceptor) (Fig. 2A,B, Supplemental table 1) emerging as the top MDH2 binders in the eCLIP analysis. Arg TCT and Glu TTC tRNA isoacceptors also feature among the cytosolic tRNAs with enriched MDH2 crosslinking sites. In addition, MDH2 binds to several other classes of cytosolic RNAs including mRNAs, lincRNAs, rRNAs and others (Fig. 2A). Highly abundant mRNAs like PABPC1 and CLU (Fig. 2B) comes up as top hits among mRNAs. We also analysed the distribution of crosslinking sites on MDH2’s previously identified interactors (SCN1A, lncRNA AC020978, GAS5) (Chen et al., 2017; Sang et al., 2021; Xu et al., 2021) in our eCLIP-dataset. Crosslinking sites of MDH2 on SCN1A mRNA were not detected in our eCLIP experiment performed in Huh7 cells, which could be due the predominantly brain-specific expression of this mRNA (Heighway et al., 2022). While we detected crosslinking sites on the other previously identified MDH2 interactors, lncRNA AC020978 and GAS5, these were not enriched over the SMI control in Huh7 cells.

To assess the binding of MDH2 to a target RNA *in vitro*, we selected one of the tRNAs enriched in the MDH2 eCLIP dataset, Arg TCT 3-2. Based on initial screening, A chemically synthesised 35 nt RNA probe matching a segment of Arg TCT 3-2 tRNA, tagged with a fluorescent Cy5 label at the 3’ end, was used to assess RNA binding. We also designed a control probe derived from the 5’ UTR of RPS23 mRNA (not enriched in the MDH2 eCLIP dataset) to evaluate the specificity of MDH2-RNA interactions. Full length human MDH2 including the N-terminal mitochondrial targeting signal (MTS) and corresponding to the cytosolic preprotein, was expressed and purified from the *Escherichia coli*-Rosetta (DE3) strain (Supplemental Fig. 3A,B,C). A C-terminal Strep-II tag was used for protein purification and retained for the binding assays. eGFP protein, purified similarly and bearing the same tag (Supplemental Fig. 3A,D), served as a specificity control.

*In vitro* electromobility shift assays (EMSA) confirmed MDH2’s interaction (0.18 μM - 5.9 μM, lanes 11-17) with the Arg TCT 3-2 probe (Fig. 2C,D) that was fixed at 10 nM in all experiments. Comparison with the control RNA probe (lanes 19-26) and the lack of binding to eGFP (lanes 1-8) confirm the specificity of the interaction (Fig. 2C,D). By quantifying the fraction of RNA bound at increasing protein concentrations, we estimated MDH2’s binding affinity to the Arg TCT 3-2 RNA probe (Kd = 0.22 μM) and to the control RNA (Kd = 1.5 μM) (Fig. 2D), suggesting that MDH2 can discriminate between different RNA fragments and confirming Arg TCT 3-2 tRNA as a preferred binding partner.

### MDH2 binds RNA preferentially outside of the mitochondria

The cytosolic MDH2 preprotein is imported into the mitochondrial matrix, where it is cleaved to form the mature MDH2 protein (Aziz et al., 1981). Considering that MDH2 associates with cytosolic RNAs, we wanted to determine the subcellular locale of MDH2’s RNA-binding. We performed comparative PNK assays on MDH2 purified from whole cell versus mitochondrially-enriched fractions (Fig. 3A). The purity of the mitochondrial fractions was confirmed by enrichment of the outer mitochondrial membrane marker, TOM20, and the depletion of cytosolic proteins (DHX9) in the mitochondrial lysates (Fig. 3B). Consistent with MDH2’s evident localisation in the mitochondrial matrix, we observed higher MDH2 signals in the input and the MDH2-immunoprecipitated samples from the mitochondrially enriched cell-fractions (Fig. 3C). Nonetheless, the PNK signal intensity for MDH2 was markedly lower from the mitochondrial fraction than from the whole cell lysate (Fig. 3C,D). Thus, combined with MDH2’s preference for cytosolic tRNAs, this result indicates that MDH2 binds RNA preferentially outside of mitochondria.

**Figure 3.**
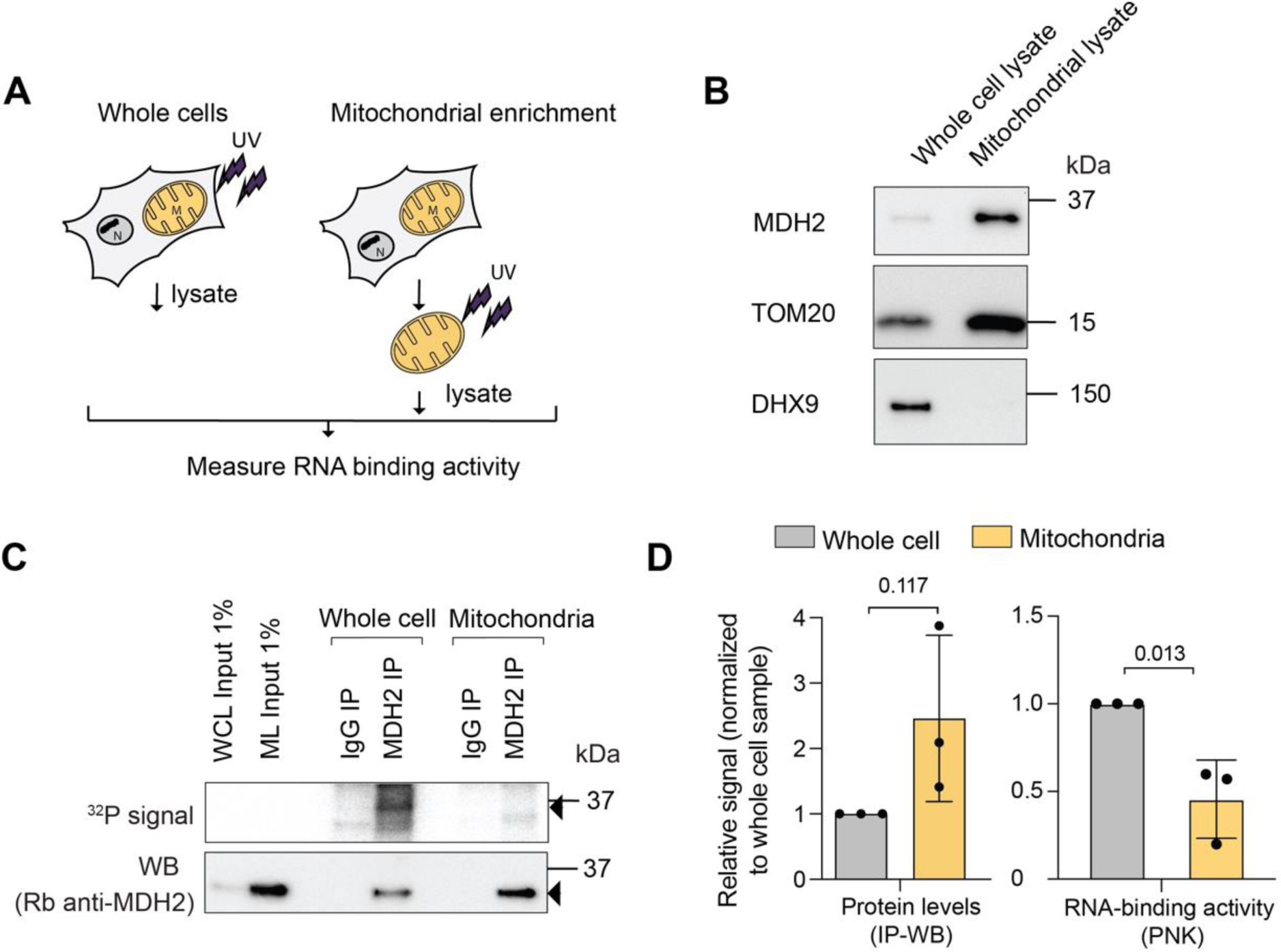
MDH2 binds RNA preferentially outside of mitochondria. (A) Experimental scheme. (B) Western blot confirming mitochondrial enrichment; TOM20, an outer mitochondrial protein, was used as a mitochondrial marker, DHX9, a cytosolic protein, was used to assess the purity of the mitochondrial prep. (C) PNK assay for MDH2 reveals a stronger RNA signal from whole cell lysates compared to mitochondrially-enriched fractions. Black arrows indicate bands near expected molecular mass of MDH2 (D) Quantification of immunoprecipitated MDH2 (IP-WB) and co-purified RNA (IP-PNK). Data are reported as mean ± S.D. number of experiments, n=3, individual p-values are indicated according to an unpaired two-tailed t-test.

To further characterize functional parameters of MDH2’s RNA-binding, we mutated MDH2 to be constrained to the cytosol. MDH2’s mitochondrial targeting signal was surmised to correspond to the N-terminal 1-24 amino acids by sequence similarity to the MDH2 targeting signals from other organisms including rat (94.4% identity with the human MDH2) (Chien et al., 1984; T. W. Chu et al., 1987; Grant et al., 1987; Sztul et al., 1988, 1989), where extensive mitochondrial import studies had been performed. This region possesses the typical characteristics of a mitochondrial targeting signal, with basic, positively charged and hydroxylated residues (Busch et al., 2023; Wiedemann & Pfanner, 2017). Based on this information, we deleted this N-terminal region from MDH2 (MDH2 Δ2-24), and evaluated the effect of this deletion on MDH2 localisation and RNA binding (Supplemental Fig. 4A).

To our surprise, deletion of amino acids 2-24 did not confine MDH2 Δ2-24 to the cytosol (Supplemental Fig. 4D). Instead, the MDH2 Δ2-24 mutant protein displays co-staining with the mitochondrial marker TOM70, indicating intact mitochondrial targeting. MDH2 Δ2-24 retained RNA binding in PNK assays, marking the canonical targeting signal as dispensable for MDH2’s RNA-binding activity (Supplemental Fig. 4B,C). Previous studies of yeast mitochondrial MDH (sharing 52.8% identity with the human mitochondrial MDH2) revealed additional internal mitochondrial localisation signals functional in the absence of a canonical presequence (Small & McAlister-Henn, 1997; Thompson & McAlister-Henn, 1989). Furthermore, cryptic targeting sequences had been observed for other yeast mitochondrial proteins like the F1-ATPase beta subunit (Bedwell et al., 1987).

Therefore, we carefully examined MDH2 for additional potential import signals using the MitoFates software (Fukasawa et al., 2015). When the full length MDH2 protein sequence was used, the expected canonical targeting signal is detected, displaying an amphiphilic alpha helical region (Fig. 4A) followed by a MPP (mitochondrial processing peptidase which generates an intermediate) cleavage site at position 16 and an Oct1 (precursor intermediate peptidase) cleavage site at position 24. When MDH2 Δ2-24 was used as the input for the MitoFates algorithm, a potential cryptic targeting signal was identified within amino acids 25-46, with an additional MPP cleavage site detected at amino acid 54. This sequence region includes a TOM20 recognition motif and a sequence stretch likely to form an amphiphilic alpha helix that could assist in the import of MDH2 Δ2-24 (Fig. 4A). Based on this information, we deleted amino acids 2-53, containing all potential import-relevant sequences of MDH2 (Fig. 4B). On analysing the subcellular localisation of MDH2 Δ2-53, we observed diffuse cytoplasmic staining in contrast to the wildtype protein, indicating deficient import into the mitochondria (Fig. 4C).

**Figure 4.**
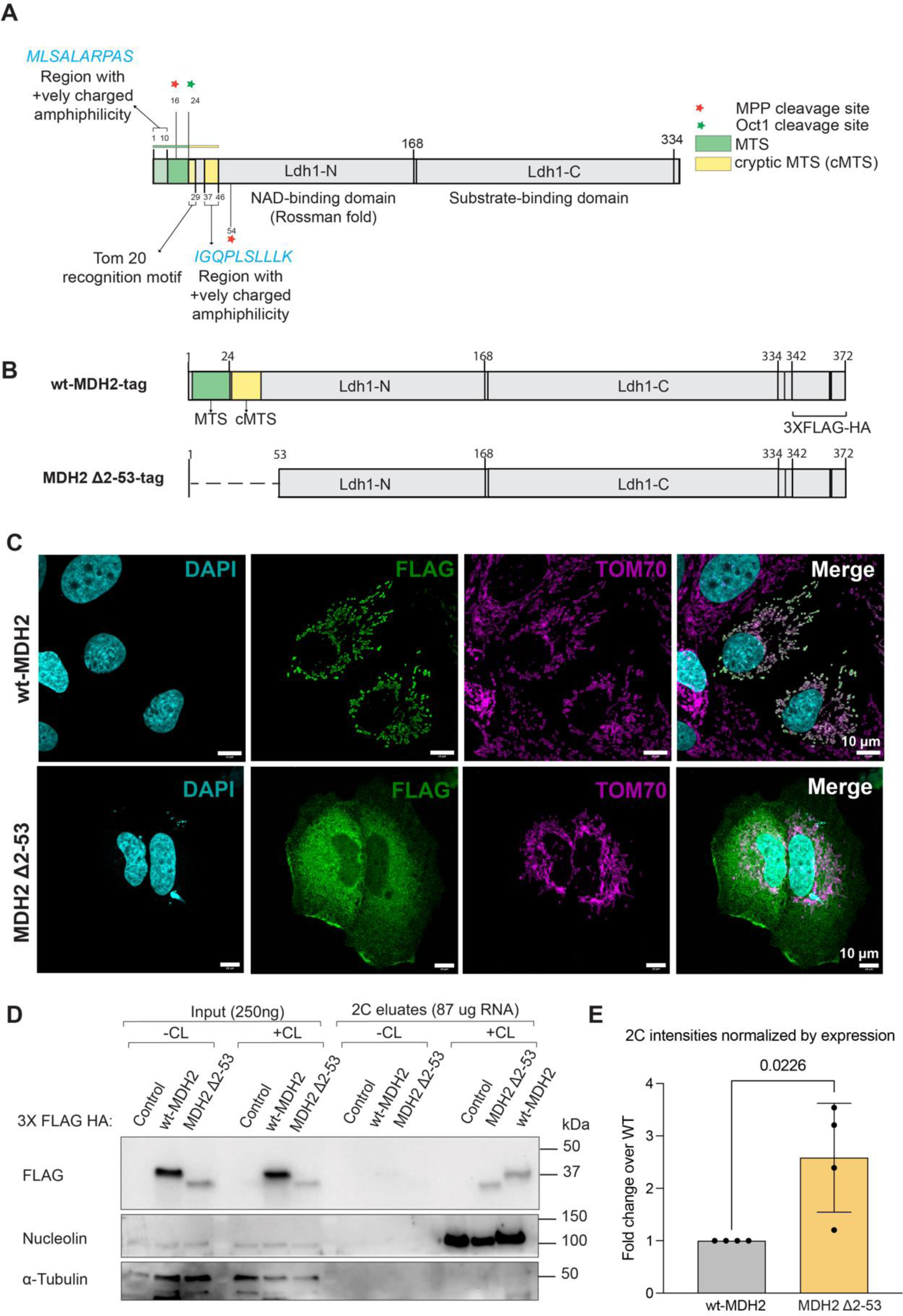
Mitochondrial targeting is dispensable for MDH2’s RNA-binding. (A) Schematic of MDH2 with the sequences relevant for mitochondrial import highlighted. MitoFates software detected the presence of amphipathic alpha helices at the N-terminus of both wt-MDH2 (highlighted in green as canonical MTS) and MDH2 Δ2-24 (highlighted in yellow as a cryptic MTS). At the N-terminus of MDH2 Δ2-24, a Tom20-recognition motif was also detected. (B) Schematic of the wt-MDH2 construct and MDH2 Δ2-53 construct. (C) Confocal immunofluorescence images showing localisation of the wt-MDH2 and MDH2 Δ2-53 construct. Images are maximum intensity projections of Z-stack images. (D) RNA-binding activity of wt-MDH2 and MDH2 Δ2-53 assessed by 2C experiments. Nucleolin is used as a positive control for RNA binding, and alpha-tubulin is used as a negative control. (E) Relative quantification of the RNA-binding activity of wt-MDH2 and MDH2 Δ2-53 normalized by the expression of the two constructs. Data reported as mean ± S.D., number of experiments, n=4, individual p-values are indicated according to an unpaired two-tailed t-test.

Next, we expressed wt-MDH2 and MDH2 Δ2-53 in Huh7 cells, and assessed RNA binding by the 2C assay. We observed lower protein expression levels for the cytosolic version of MDH2 compared to the wt-MDH2 (Fig. 4D), possibly due to a lower stability of the protein. Nevertheless, we found that MDH2 Δ2-53 retains similar or even elevated RNA binding (Fig. 4D) compared to wt-MDH2 after normalisation for the differences in expression levels (Fig. 4E). This finding reveals that mitochondrial targeting is not required for MDH2’s RNA-binding. We also conclude that the N-terminal 53 amino acid sequences are dispensable for MDH2’s RNA interactions.

### MDH2’s RNA-binding is sensitive to cellular NAD^+^ levels

Next, we wanted to explore whether MDH2’s RNA-binding responds to cellular signals. Since MDH2 binds NAD^+^ as a cofactor, and NAD^+^ levels have been reported to affect the RNA-binding properties of other dehydrogenases (Pioli et al., 2002; Rodriguez-Pascual et al., 2008; Singh & Green, 1993; Sioud & Jespersen, 1996) we sought to modulate cellular NAD^+^ levels and to monitor the effects on MDH2’s RNA-binding activity in cells. FK866, an inhibitor of the rate-limiting step of the cellular salvage NAD^+^ synthesis pathway (Khan et al., 2006; Tan et al., 2013), was used to reduce cellular NAD^+^ cells. We also supplemented cells with the NAD^+^ precursor NAM (Garten et al., 2009) to boost cellular NAD^+^ levels (Fig. 5A,B). Profound reduction of cellular NAD^+^ levels by FK866 treatment significantly activated MDH2’s RNA-binding (Fig. 5C,D). By contrast, NAM treatment yielding a 1.5-fold increase in cellular NAD^+^ levels did not substantially change MDH2’s RNA-binding activity (Fig. 5C,D). The sensitivity of MDH2-RNA interactions to FK866 treatment suggests that NAD^+^ levels may directly or indirectly control MDH2’s RNA-binding.

**Figure 5.**
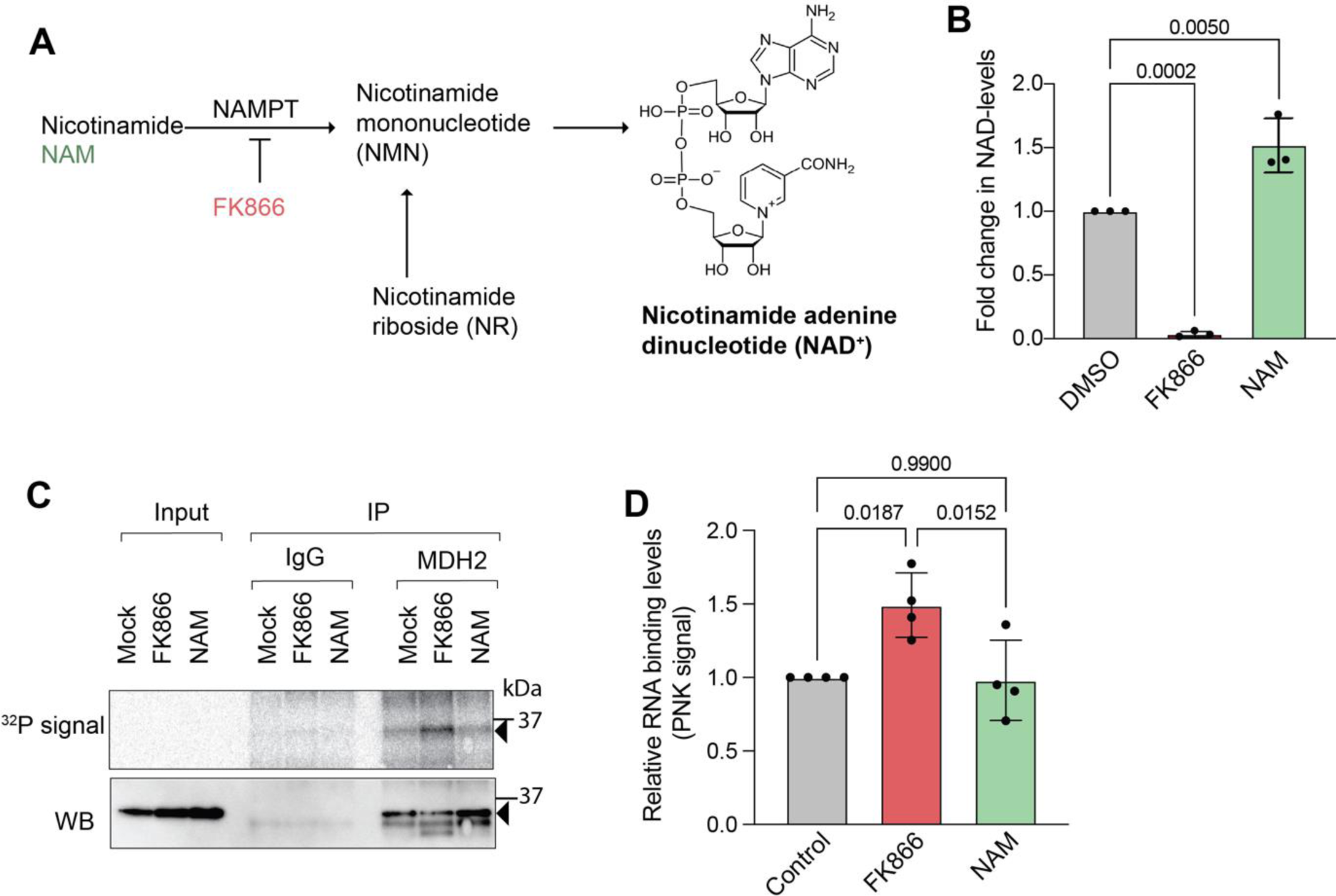
Pharmacological perturbation of cellular NAD+ levels affects MDH2’s RNA-binding. (A) Scheme of the salvage synthesis pathway for NAD+ showing the steps targeted by NAM and FK866 treatment, respectively. (B) Changes in cellular NAD(H) levels measured after 48 h of treatment. (C) PNK assay to assess MDH2’s RNA-binding on alteration of cellular NAD(H) levels. Black arrows indicate expected molecular mass of mature MDH2. (D) Quantification of RNA binding (PNK signal) of MDH2 after NAM/FK866 treatment. Data reported as mean ± S.D, number of experiments, n=3, individual p values as calculated by ordinary one-way ANOVA with Tukey’s multiple comparison test correction.

## Discussion

In this study, we have characterised the RNA binding of the human mitochondrial enzyme MDH2. We confirmed cellular RNA binding using two orthogonal assays, and identified its RNA interactors by eCLIP. tRNAs, in particular, emerged as top binders, being significantly enriched in IP compared to SMI controls. Whether MDH2 binds full length tRNAs or tRNA fragments that have been described to be generated especially under conditions of cellular stress (Su et al., 2020; Xie et al., 2020) remains to be defined. MDH2’s preference for binding tRNAs is also supported by studies in yeast, which identified yeast MDH2 as an RBP that predominantly binds to small RNAs (Asencio et al., 2023). Interestingly, tRNA-binding has also been reported previously for the NAD-dependent dehydrogenase GAPDH (Nagy & Rigby, 1995; Singh & Green, 1993).

On investigation of the subcellular location of MDH2-RNA interactions by mitochondrial purification followed by RNA-binding assays, we found that MDH2 binds RNA predominantly outside of mitochondria, which is in excellent accord with the finding that cytosolic RNAs represent the major class of interactors. We also determined that MDH2’s RNA-binding does not require the N-terminal 52 amino acids of the precursor protein or its mitochondrial localisation/targeting. The former implies that the mature protein that reaches the mitochondria bears all amino acids required for RNA binding, although this major MDH2 fraction does not contribute substantially to the overall amount of RNA bound by MDH2 in the conditions we have assayed. The latter observation is in further support of the predominantly cytosolic nature of the MDH2-RNA interaction. We envision that RNA binding by MDH2 may be determined by its environment, which is discussed in more detail below.

Primarily cytosolic RNA binding of a mitochondrial metabolic enzyme is somewhat unexpected, and there are limited data from other systems (Wang et al., 2017). Following viral infection, mitochondrial glutamic-oxaloacetic transaminase (GOT2) binds lncRNA-ACOD1, with an increased fraction of GOT2 appearing in the cytosol. The expression of lncRNA-ACOD1 is induced by viral infection, but it is currently not clear whether the RNA-binding pool of GOT2 is predominantly the precursor form or the mature GOT2 protein present in the cytosol by an as yet-unidentified mechanism. With the exception of proteins like cytochrome c that are released into the cytosol from mitochondria during conditions of cellular stress (Garrido et al., 2006), there is little evidence of mature mitochondrial proteins being released back into the cytosol. There have been reports of mitochondrial TCA cycle enzymes (Kafkia et al., 2022; Nagaraj et al., 2017) and the mitochondrial pyruvate dehydrogenase complex being localised to the nucleus (Sutendra et al., 2014). Translocation across the mitochondrial membranes, followed by nuclear export and mitochondria-derived vesicles (Sugiura et al., 2014), has been discussed as potential mechanisms of a cytosolic localisation in cases where nuclear localisation signals were absent. Examination of the MDH2 signal from PNK and 2C assays does not allow a clear assignment of whether the MDH2 preprotein or mature MDH2 binds RNA. The molecular mass difference between the two is only ∼2-3 kDa, and the effect of crosslinked RNA on gel migration must also be taken into account. Therefore, both possibilities remain open for now.

At present, possible functional roles of the MDH2-RNA interactions include both “moonlighting” of MDH2 as a posttranscriptional effector of the fates of the bound RNAs as well as riboregulation of MDH2 function by the bound RNAs (Hentze et al., 2018). While a “moonlighting” role has been suggested in a previous study (Chen et al., 2017), a rigorous analysis using carefully designed RNA-binding mutants is needed to deconvolute the situation. In addition to previously described effects of RNA on TCA cycle activity by disrupting a FH-MDH2-CS metabolon (Sang et al., 2021), other regulatory effects of RNA on MDH2 can also be imagined, including direct modulation of MDH2’s enzymatic activity, regulation of its mitochondrial function by cytoplasmic sequestration, or modulation of mitochondrial import.

Our finding that MDH2-RNA interactions respond to cellular NAD^+^ levels may indicate physiological regulation by the concentration of this cofactor and/or other related metabolites. Nucleotide binding domains, especially NAD-binding domains, have been associated with RNA binding of metabolic enzymes in several RNA interactome captures (Castello et al., 2015; Perez-Perri et al., 2023). For RNA-binding dehydrogenases like GAPDH (Rodriguez-Pascual et al., 2008; Singh & Green, 1993; Sioud & Jespersen, 1996) and LDH (Pioli et al., 2002), NAD^+^ has been reported to compete with RNA binding. Thus, NAD^+^ may also compete with RNA to bind MDH2, directly or allosterically, explaining the observed increase in MDH2’s RNA-binding following FK866 treatment (Fig. 5). Extending this hypothesis further, compartmentalised NAD^+^ levels in the cytosol and mitochondria could be a determinant of the compartmentalisation of MDH2’s RNA-binding, the typically lower concentration of NAD^+^ in the cytosol being less restrictive to RNA binding than the higher mitochondrial NAD^+^ levels (Alano et al., 2007; Cambronne et al., 2016; Di Lisa et al., 2001; Sallin et al., 2018). However, precise quantification of NAD^+^ levels across compartments is challenging, and there are limited data available on the absolute concentration of NAD^+^ in cytosol compared to the mitochondria (VanLinden et al., 2015). Moreover, MDH2 activity has been found to be regulated by acetylation, a conceivable indirect effect of NAD^+^ modulation via NAD^+^-dependent SIRT deacetylases (Marcus & Andrabi, 2018; Parodi-Rullan et al., 2018). Therefore, acetylation-driven regulation of MDH2’s RNA-binding also deserves to be considered.

In conclusion, our data highlight an unexpected facet of the RNA-binding activity of a mitochondrial enzyme which is not yet functionally understood, but that serves as an example of the scope of RNA-protein interactions in cells. The strong conservation of RNA binding by MDH2 and the effect of altered NAD^+^ levels on RNA binding suggest that the RNA-binding properties of MDH2 warrant further exploration.

## Materials and Methods

### Cell culture

Huh7 cells were maintained in a 37°C incubator with 5% CO2, with Dulbecco’s modified Eagle’s medium (DMEM) growth media (reconstituted from Sigma D5523) supplemented with supplemented with 1 g/L D-glucose (Sigma), 10% heat inactivated FBS (Thermo Fischer Scientific 10270106), 2 mM L-glutamine Thermo Fischer Scientific 25030081), and 100 U/ml Penicillin-streptomycin (Thermo Fischer Scientific 15140122).

### Protein extracts, SDS-PAGE and western blotting

For preparation of cell lysates, cells were washed twice with ice-cold PBS and lysed in RIPA buffer (50 mM Tris/HCl, pH 7.5, 150 mM NaCl, 1% NP40, 0.1% SDS, 0.5% Na-deoxycholate), supplemented with fresh complete protease inhibitor cocktail. The lysates were sonicated and cleared by centrifugation at 10000 g for 15 min. Protein amounts were estimated with Pierce 660 reagent (Thermo Fischer Scientific 22660) against defined BSA standards (Thermo Fischer Scientific 23208). Samples were prepared for electrophoresis by addition of 4x loading buffer (Thermo Fischer Scientific NP0007) supplemented with TCEP as a reducing agent and incubated at 70°C for 10 min. 4-15% Tris-Glycine (BioRAD precast gradient gels 5671084) were used for protein separation. Proteins were transferred to PVDF/nitrocellulose membranes using the Transblot semi-dry blotting system (Bio-Rad). Transfer was usually perfomed at 25 V for 7 min. The membranes were incubated with the blocking buffer (5% milk in 0.05% PBST) for 30 min to 1 h at room temperature. This was followed by incubation with primary antibodies against the target protein at 4°C O/N or 1-3 h at room temperature on a rotator. The primary antibody was diluted in the blocking buffer. The membrane was washed 3 times with 0.05% PBST with 5-minute incubations. After washing off the unbound primary antibody, the membrane was incubated at room temperature (RT) for 30 min in HRP-conjugated secondary antibody diluted in the blocking buffer. After 3 washes with 0.05% PBST with 5-minute incubations, the membranes were developed in ECL solution (Sigma WBKLS0500) or the SuperSignal West Femto/Atto solutions (Thermo Fisher Scientific A37558, 34095). The antibodies used were: rabbit polyclonal MDH2 antibody (Proteintech 15462-1-AP, 1:1000), goat MDH2 antibody (Everest Bio EB13027, 1:1000), FLAG (Sigma-Aldrich F1804, 1:2000) DHX9 (Abcam ab26271, 1:5000), Tom20 (Proteintech 11802-1-AP, 1:1000), Tom70 (Proteintech 14528-1-AP, 1:1000), Nucleolin (Abcam ab50279, 1:1000), HuR (Proteintech 11910-1-AP, 1:1000).

### PNK assay

Cells (80-90% confluency) were washed twice with ice-cold PBS and irradiated with UV light (254 nm) of 150 mJ/cm2 energy on ice. The cells were lysed in RIPA buffer. The lysates were sonicated and cleared by centrifugation at 10000 g for 15 min at 4 °C. Around 200-600 μg of total protein lysate was used for PNKs. The lysates were treated with 0.1-0.6 ng/μl RNaseA (Thermo Fischer Scientific EN0531) and 2U of Turbo DNAse (Thermo Fischer Scientific AM2239) for 15 min at 37°C. The lysates were spun down briefly and a fraction of lysate set aside as input to assess IP efficiency in a western blot. The remaining lysates were for immunoprecipitations. For immunoprecipitation of MDH2, SureBeads ProteinA magnetic beads (Bio-Rad 1614013) coupled with MDH2 antibody (Proteintech 15462-1-AP) were used. An isotype matched rabbit IgG (Cell Signalling 02/2018) coupled to SureBeads ProteinA magnetic beads were used in parallel immunoprecipitations as background control. For immunoprecipitation of overexpressed FLAG-tagged MDH2 variants, FLAG M2 beads (Merck M8823) were used. The lysate-antibody coupled bead mixture was incubated at 4°C for 2 hours. The beads were washed with RIPA buffer (50 mM Tris/HCl, pH 7.5, 150 mM NaCl, 1% NP40, 0.1% SDS, 0.5% Na-deoxycholate, fresh complete protease inhibitor cocktail), RIPA-HS buffer (50 mM Tris/HCl, pH 7.5, 500 mM NaCl, 1% NP40, 0.1% SDS, 0.5% Na-deoxycholate, fresh complete protease inhibitor cocktail) and PNK buffer (50 mM Tris/HCl, pH 7.5, 50 mM NaCl, 0.5% NP40, 10mM MgCl2, fresh 5mM DTT, fresh complete protease inhibitor cocktail) for two minutes each. For labelling the RNA, the beads were incubated in the 30 μl T4 PNK reaction mix (0.3 μl of 0.1 μCi/μl γ-32P ATP, 3U of T4 PNK enzyme (NEB M0201L) in 1X PNK reaction buffer containing 1mM DTT) for 15 min at 37°C. Following 4 washes with the PNK buffer, proteins were eluted off the beads at low pH (0.1 M glycine, pH 2.0) for endogenous MDH2 IP and incubation with 250 μg/ml 3XFLAG peptide (Merck F3290) in the case of FLAG IP. When eluted at low pH, the sample solution was neutralised by the addition of 1.5 M Tris-HCl, pH 8.5. After addition of 4x sample loading buffer (Thermo Fischer Scientific NP0007) with added TCEP (Merck 646547), the samples were boiled for 10 min at 70°C. The samples were resolved using SDS-PAGE and further transferred to a nitrocellulose membrane. After drying the membrane, it was exposed to a phosphor screen overnight. The screen was scanned on a Typhoon FLA 9500 system at 600 nm. The membrane was further immunoblotted with MDH2 antibody to estimate IP efficiency.

### 2C assay

Complex capture/2C was performed as described (Asencio et al., 2018). Huh7 cells were grown to 80-90% confluency in 15cm dishes. Cells were washed twice with ice-cold PBS. and irradiated with UV light (254 nm) of 150 mJ/cm2 energy on ice. The cells were lysed in RIPA buffer (50 mM Tris/HCl, pH 7.5, 150 mM NaCl, 1% NP40, 0.1% SDS, 0.5% Na-deoxycholate, fresh complete protease inhibitor cocktail). The lysates were sonicated and cleared by centrifugation at 10000 g for 15 min. 1-1.5 mg of samples were processed for 2C. A fraction of the input was set aside for western blot assessment. Samples were processed according to the manual using the components of the Zymo RNA mini kit (Zymo R1013). The RNA and RNA-protein complexes were eluted from the column with 100 uL Nuclease free water. The eluates were treated with 5U Turbo DNAse (Thermo Fischer Scientific AM2239) for 30 min at 37°C. In selected samples, 5 U RNase I (Ambion AM2294) was included in the reaction. The samples were then processed for a second round of complex capture as described above. The eluates were treated with RNase I (Ambion AM2294) (with exceptions) to digest the eluted RNA in the RNP complexes and visualise the protein around the expected molecular weight using western blotting. Samples were mixed with 4x sample loading buffer (Thermo Fischer Scientific NP0007) supplemented with TCEP and incubated at 70°C for 10 min before gel-loading. The samples were resolved using SDS-PAGE and immunoblotted with the antibodies of interest.

### eCLIP

eCLIP was performed according to published protocols (Van Nostrand et al., 2016) with the following changes. 75 μl of SureBeads ProteinA magnetic beads (Bio-Rad 1614013) were coupled with 6 μg of MDH2 antibody (Proteintech 15462-1-AP) for 2 h at 4°C. 500 μg precleared cell protein lysate was treated with 1 U RNase I (Ambion AM2294) for 5 min at 37°C. After addition of 4 μl of RNAsin Plus ribonuclease inhibitor (Promega N2611), the samples were spun down, transferred to a new tube and used for immunoprecipitation (IP) with the antibody coupled beads. Before IP, 1% of the input was set aside as size-matched input control. The IP was performed at 4°C for 2 h. Isotype matched IgG was used as a control. Following library preparation till the cDNA step as described previously (Van Nostrand et al., 2016), we optimised the number of cycles required for the final PCR using test PCRs with 10% of the cDNA sample followed by gel analysis (6% TBE gels, Thermo Fischer Scientific EC6265BOX). cDNA libraries from MDH2 IP (14 cycles), IgG IP (14 cycles) and SMI (9 cycles) were multiplexed and analysed with paired end sequencing on NextSeq 500 using different cycle numbers for read 1 (20 cycles) and read 2 (130 cycles), with index 1 and 2 read by 8 cycles each.

### eCLIP data analysis

The quality of the eCLIP raw reads was examined using fastqc (Andrews, 2010) (v0.11.8). The Unique molecular identifier (UMI) barcodes, attached during library preparation, were appended to the read name using UMI tools (v1.0.0) extract (Smith et al., 2017). The adapters were trimmed and shorter reads (length < 18 nucleotides) were discarded using cutadapt (v2.5) (Martin, 2011). The trimmed reads were mapped to the human genome (GRCh38.v23 from GENCODE) using STAR(v2.7.1a) (Dobin et al., 2013). PCR duplicated reads from the aligned reads were removed using UMI tools dedup based on the UMI barcode.

The tRNAs, provided by tRNAscan (Lowe & Chan, 2016), were added to the GENCODE (v23) annotation of the GRCh38.v23 genome and preprocessed with the htseq-clip suite (Sahadevan et al., 2022). These annotations were re-formatted into sliding windows of 50 nucleotides with a step size of 20 nucleotides, using htseq-clip annotation and createSlidingWindows functions. Crosslink site was extracted as one nucleotide upstream of the alignment start position for each aligned read using the htseq-clip extract function. htseq-clip count was used to quantify the total number of crosslink sites per sliding window and was converted into an R-friendly count matrix using htseq-clip createMatrix function. The count matrix was prefiltered to retain only those windows with a minimum of 10 cross-link sites in at least 3 samples. DEWSeq (v1.0.0) (Schwarzl et al., 2023), a R/Bioconductor package, was used to determine significant crosslink enrichment of windows in IP samples over the size-matched input (SMI) control samples (p_adj_ ≤0.05 and log_2_FC≥ 1). The Bonferroni method (Bland & Altman, 1995) was used to control the familywise error rate, and the Benjamini– Hochberg method (Benjamini & Hochberg, 1995) was used to control the false discovery rate. Windows with significant enrichment in MDH2 IP in comparison to SMI (p_adj_ ≤0.05 and log_2_FC≥ 1) were selected for further downstream analysis and overlapping windows among these were merged into binding regions. These binding regions (782) were then screened to remove those binding sites with zero read counts in the input samples (to prevent artificial enrichment), as well as aberrant single-peak crosslinking sites (mostly in intronic regions). To generate Fig. 2A, the number of crosslinking counts of MDH2 on the identified regions (aggregated normalized counts of crosslink-sites across all windows in the region) was used (Supplemental Table 1).

### Protein expression and purification

petM22 protein purification constructs were transformed into BL21 Rosetta (DE3) cells. The transformed bacteria were grown overnight at 37°C in 25 ml Lysogeny broth (LB) media supplemented with Kanamycin (Sigma K0254). For autoinduction of MDH2 and eGFP proteins, the preculture was diluted 1:100 in Terrific Broth (TB) supplemented with 1.5% lactose, 0.05 % glucose, 2 mM MgSO4 for growth at 37°C till the OD reached 0.7-0.8. The bacterial culture was then transferred to an 18°C incubator for overnight growth with shaking at 180 rpm. The Strep-Tactin Superflow high-capacity resin (IBA lifesciences, 2-1208-002) was used to purify the protein. First, the cells were lysed by sonication in 100 mM Tris/HCl, pH 8.0, 150 mM NaCl, 1 mM EDTA supplemented with free protease inhibitors and 250 μg/ml lysozyme. The cells were sonicated for 7 minutes at 50% duty, 70% power on wet ice and centrifuged at 15000 g for 15 min to remove debris. The supernatant was allowed to flow through a polypropylene column packed with Strep-Tactin resin. The column was washed five times with 100 mM Tris/HCl, pH 8.0, 150 mM NaCl. The column-bound protein was eluted in fractions with 100 mM Tris/HCl, pH 8.0, 150 mM NaCl, 1 mM EDTA, 2.5 mM desthiobiotin (IBA lifesciences, 2-1000-002). The eluted fractions containing highest concentration of the expressed protein were pooled and dialyzed against the storage buffer containing 10mM Tris/HCl, 150mM NaCl, 5% Glycerol. The sample concentration was estimated by UV-Vis spectroscopy (absorbance at 280 nm) and using defined BSA standards (Thermo Fischer Scientific 23208) through SDS-PAGE analysis. If needed, the samples were concentrated using Amicon Ultra-0.5 Centrifugal Filter Unit (Millipore, UFC5010). The purified protein aliquots were stored at -80°C.

### Electromobility shift experiments

Binding reactions containing 10 nM Cy5-labelled RNAs (IDT, Sigma) and defined amounts of proteins were set up in 10 mM Tris-HCl pH 7.5, 150 mM NaCl, 20% glycerol, 0.5 mM DTT, 5 mM MgCl2, 0.01 μg/ul BSA and incubated at 25°C for 20 minutes. The probe sequences for the Arg TCT 3-2 tRNA and the control probe were respectively GGCUCUGUGGCGCAAUGGAUAGCGCAUUGG and AUAUUUAGCUCAGGCCCGUGGCGCC. The binding reactions were resolved and analysed using native polyacrylamide gel electrophoresis (4-15% Tris-Glycine gradient gels in running buffer containing 34 mM Tris-HCl, 66 mM HEPES, 0.1 mM EDTA), with the separation done at 100V for 75 min. Fluorescent signal on the RNA was analysed using Typhoon FLA-9500.

### Mitochondrial fractionation and crosslinking

Cells grown to 80-90% confluency were washed two times with ice-cold PBS and harvested in 3 ml of 20 mM HEPES pH 7.4-7.6, 220 mM Mannitol, 70 mM Sucrose, 1 mM EDTA supplemented with 2 mg/ml BSA, complete EDTA-free protease inhibitors and RNAsin (Promega N2611). Cells and allowed to swell by incubation at 4°C for 10 min and lysed with 20-25 strokes of a Dounce homogenizer. Nuclei was pelleted by centrifugation at 1000 g for 10 min at 4°C. The supernatant was centrifuged at 15000 g for 20 minutes at 4°C to pellet mitochondria. The mitochondrial pellet was washed 2 times with 20 mM HEPES pH 7.4-7.6, 220 mM Mannitol, 70 mM Sucrose, 1 mM EDTA. The wash buffer was removed by centrifugation for 20 minutes at 4°C. The pellet was resuspended in 100 μl of 20 mM HEPES pH 7.4-7.6, 220 mM Mannitol, 70 mM Sucrose, 1 mM EDTA and then irradiated with UV light after spreading on a glass bottom dish. The mitochondria were lysed by sonication after adding excess volume of RIPA buffer (50 mM Tris/HCl, pH 7.5, 150 mM NaCl, 1% NP40, 0.1% SDS, 0.5% Na-deoxycholate, fresh complete protease inhibitor cocktail). The lysates were snap frozen and stored at -80°C till further use. Whole cell samples were processed at the same time by crosslinking and lysis in RIPA buffer (50 mM Tris/HCl, pH 7.5, 150 mM NaCl, 1% NP40, 0.1% SDS, 0.5% Na-deoxycholate, fresh complete protease inhibitor cocktail).

### Transfections

Cells were transfected using either the forward (plating cells on the day before) or reverse method (plating cells right before) transfection. Lipofectamine 3000 reagent (Themo Fischer Scientific 11668030) was used according to the manufacturer’s instructions to transfect plasmids.

### Immunofluorescence

Cells were grown to 60% confluency on coverslips/multiwell glass bottom dishes (ibidi) and fixed with 4% paraformaldehyde (Themo Fischer Scientific 28906). 0.2% Triton-X100 was used to permeabilize cells by incubation at room temperature for 10 minutes. Samples were blocked in the IF blocking buffer (2.5% BSA in 0.05% TritonX100) for 30 minutes at room temperature. After blocking, samples were incubated with antibody diluted in IF blocking buffer for 1 hour at 37°C. Samples were washed 3 times in 0.05% TritonX100 before incubating with the secondary antibody labelled with the fluorescent dye (also diluted in the blocking buffer) for 45 min to 1 hour at room temperature in the dark. The following secondary antibodies from Life technologies were used: anti-rabbit Alexa Fluor 488 (A21206), anti-mouse Alexa Fluor 488 (A21202), anti-rabbit Alexa Fluor 647 (A21245). Nuclei were stained by incubation with Nucblue reagent (Themo Fischer Scientific R37606) diluted in 0.05% TritonX100 (Invitrogen) for 10 min. The samples were further washed two times before mounting on a glass slide using ProLong Diamond antifade mountant (Invitrogen P36961) or addition of mounting solution (ibidi 50001) in glass dishes. The glass slides were left at room temperature to cure overnight before imaging and stored for long term at 4°C. Images were acquired on an Olympus 3000FV microscope with either a 60X objective/ 40X objective.

### Cellular NAD modulation

Cells growing at around 60% confluency were treated with 5 nM FK866 (Merck 481908), 5 mM NAM (Sigma N0636) or DMSO (Merck 1.02950) for 48 hours before harvest. A fraction of the cell lysates was used to estimate NAD(H) levels using the NAD/NADH quantification kit (Sigma MAK037) according to the manufacturer’s instructions.

## Data availability

Sequencing data has been submitted to GEO and is available under the following accession number: GSE249145.

## Quantification and Statistical Analysis

GraphPad Prism version 9.3.1 was used for statistical analysis of experimental data (excluding sequencing data). The tests used are documented in the figure legends wherever applicable.

## Acknowledgements

We thank current and former members of the Hentze laboratory for helpful suggestions. We thank Sudeep Sahadevan for helping with the data analysis and input on the manuscript. We acknowledge EMBL’s core facilities, specifically the genomics, protein expression and purification, advanced light microscopy core facilities for their guidance and expert services. We gratefully acknowledge insightful discussions with Dr. Anna Mandinova (Harvard Medical School and Cutaneous Biology Research Center, Massachusetts General Hospital). M.W.H gratefully acknowledges funding from Manfred Lautenschläger Foundation (Heidelberg).

## Author contributions

Conceptualization, M.N, A.C, M.W.H; Investigation, M.N, A.C; Formal data analysis and interpretation, M.N, A.C, T.Sekaran, T.Schwarzl, M.W.H; Project administration, M.N, A.C, M.W.H; Mentoring and supervision, A.C., M.W.H; Writing (original draft and editing), M.N., M.W.H. with contributions from all co-authors.

